# Potassium channels mediate the inhibitory effect of exosomes on anti-tumor immunity in head and neck cancer

**DOI:** 10.1101/2025.07.25.666838

**Authors:** Ameet A. Chimote, Abdulaziz O. Alshwimi, Simran Venkatraman, Jay Bhati, Maria A. Lehn, Somchai Chutipongtanate, Sarmistha Das, Susan Kasper, Shesh N. Rai, Scott M. Langevin, Trisha M. Wise-Draper, Laura Conforti

## Abstract

Head and neck squamous cell carcinomas (HNSCC) are aggressive cancers with a relatively low response rate to immunotherapy. Anti-tumor immune responses rely on cytotoxic T and NK cells infiltrating the tumor microenvironment to eliminate cancer cells. However, tumors activate multiple mechanisms to evade these responses. Tumor-derived small extracellular vesicles, also known as exosomes, reshape the tumor microenvironment by impairing cytotoxic T and NK cell function, promoting immune escape and metastasis. Transcriptomic analysis of healthy donor peripheral blood mononuclear cells (PBMCs) exposed to exosomes from HPV-negative HNSCC patient primary cancer cells revealed reduced cytotoxic cells and suppressed immune responses including cytotoxicity, chemokine production, and NK and T cell functions. Bead-based multiplex immunoassay showed that HNSCC-derived exosomes inhibited the release of effector cytokines (IL-2, TNF-α, IFN-γ) and cytotoxic molecules from activated CD8⁺ T cells. While ion channels regulate Ca²⁺-dependent cytotoxicity and cytokine production and release, their role in exosome-mediated immune suppression is unexplored. We found that tumor-derived exosomes selectively inhibit KCa3.1 channel activity in CD8⁺ T cells by downregulating calmodulin, ultimately impairing Ca²⁺ signaling and IFN-γ release. This study identifies a novel mechanism of exosome-mediated immunosuppression, positioning KCa3.1 as a promising therapeutic target to enhance immune surveillance and immunotherapy response in HNSCC.

## Introduction

Head and neck squamous cell carcinoma (HNSCC) is an aggressive cancer with a 5-year overall survival rate of less than 30% for high-risk cases and limited response to immunotherapy (1). Solid malignancies, including HNSCC, depend on the infiltration and cytotoxic activity of T cells to eliminate cancer cells. However, the tumor microenvironment (TME) is enriched with immunosuppressive factors, such as adenosine and inhibitory checkpoint ligands, which impair the recruitment and function of these immune cells, posing a major obstacle to effective cancer therapy (2). Targeting these immunosuppressive pathways is a key focus of clinical research— PD-1 blockade is FDA-approved for recurrent metastatic HNSCC and resectable locally advanced HNSCC expressing PD1 ligand (PDL1), and inhibitors of adenosine and TIGIT blockade have demonstrated antitumor activity in preclinical studies, particularly when combined with PD1 inhibitors (3–6).

Small extracellular vesicles, commonly referred to as exosomes, have recently been recognized as key contributors to the failure of immune surveillance in cancer. These nanoscale (30–150 nm) membrane-encapsulated vesicles, which are released by both non-pathologic and pathologic cells, including cancer cells, facilitate intercellular communication by transferring proteins and genetic material from their cell of origin (6). Tumor-derived exosomes carry immunosuppressive molecules, including PD1 ligands, CD39 and CD73 (enzymes responsible for adenosine production), and nucleic acids that alter recipient cells (6). Exosomes from HNSCC cell lines and patient sera have been shown to induce T cell apoptosis, reduce T cell infiltration and cytotoxicity in tumors, thereby promoting tumor progression in HNSCC mouse models (7–10). PDL1-positive exosomes contribute to systemic immunosuppression in mice and correlate with disease progression in HNSCC patients (9,11). Additionally, the presence of tumor-derived exosomes in patient serum is associated with metastasis as they shape the metastatic niche by activating pro-tumor signaling in stromal and endothelial cells while suppressing immune surveillance (2,12). Despite these findings, the mechanisms underlying exosome-mediated immunosuppression remain poorly understood, and the contribution of ion channels in immune cells to exosome-induced immunosuppression has yet to be explored.

T cells express multiple ion channels that regulate their effector functions (13). Kv1.3 and KCa3.1 potassium channels help maintain membrane resting potential and facilitate Ca^2+^ influx through Ca^2+^ channels (CRAC and TRPM7), which are essential for downstream functions (13). Kv1.3, KCa3.1, and CRAC channels play crucial roles in T cell activation, as their inhibition suppresses cytokine production and proliferation. Additionally, KCa3.1 and TRPM7 regulate T cell motility (14).

KCa3.1 channels also mediate the immunosuppressive effects of various TME elements. Adenosine, prostaglandin E2 (PGE2), PDL1, and nutrient deprivation within the TME have all been shown to inhibit KCa3.1 (15–18). Consistent with these findings, we have observed reduced KCa3.1 expression in tumor-infiltrating and circulating cytotoxic T cells from HNSCC patients, leading to reduced tumor infiltration and impaired cytokine production (19). This deficiency in KCa3.1 in peripheral T cells may also contribute to increased infection susceptibility and diminished vaccine responses in cancer patients (20). Despite the well-established role of ion channels in T cells, it remains unclear how the tumor signals systemic defects in KCa3.1 on peripheral T cells. Here, we provide evidence that HNSCC exosomes suppress KCa3.1 activity in T cells, facilitating tumor immune evasion. These findings uncover a novel mechanism of cancer immune escape and could lead to novel immunotherapies targeting KCa3.1 channels.

## Materials and Methods

### Human subjects

This study utilizes primary cancer cells from HNSCC patients. The characteristics of these patients have been reported in our previous publication (21). Briefly, these patients had biopsy-confirmed oral cavity squamous cell carcinoma that were tested negative for HPV and positive for PDL1, and had no prior radiation or chemotherapy. Informed consent was obtained from all participants, and the study protocol was approved by the University of Cincinnati Institutional Review Board (IRB No. 2014-4755).

### Cell culture

#### Reagents and chemicals

Dulbecco’s Modified Eagle Medium (DMEM), Roswell Park Memorial Institute Medium 1640 (RPMI 1640), a 1:1 mixture of DMEM and Ham’s F12 Nutrient Mix (DMEM/F12), phosphate-buffered saline (PBS), penicillin–streptomycin (100×), trypsin-EDTA solutions (0.25% and 0.05%), fetal bovine serum (FBS), exosome-depleted fetal bovine serum, minimum essential amino acids (MEM AA), sodium pyruvate, L-glutamine, epidermal growth factor (EGF), and insulin–transferrin–selenium supplement (ITS-G) were obtained from Gibco (Thermo Fisher Scientific, Waltham, MA, USA). Adenine, hydrocortisone, and high-purity cholera toxin were purchased from MilliporeSigma (Burlington, MA, USA). Rho-associated protein kinase (ROCK) inhibitor and Primocin antibiotic mixture were purchased from Cayman Chemical (Ann Arbor, MI, USA) and InvivoGen (San Diego, CA, USA), respectively.

#### Primary HNSCC patient-derived keratinocyte cultures

Primary keratinocyte cultures were established from HNSCC tumor tissues from three de-identified patients: HNC208, HNC285, and HNC365, as described previously (21). Briefly, the tumors were minced and enzymatically dissociated using 0.25% trypsin, then plated onto irradiated (60Gy) NIH/3T3 feeder cells in keratinocyte medium (DMEM/F12 supplemented with adenine, amino acids, sodium pyruvate, ITS-G, Primocin, hydrocortisone, cholera toxin, ROCK inhibitor, EGF, and 5% FBS). Cultures were maintained at 37°C and passaged through differential trypsinization. Mycoplasma testing was performed routinely using MycoStrip® mycoplasma detection kit (Invivogen, Catalog# rep-mys-50) and was negative for all cultures. For the final two passages before exosome isolation, cells were cultured in media with 5% exosome-depleted FBS and no feeder cells.

### Exosome Isolation and Characterization

#### Exosome isolation

Exosomes were isolated from conditioned media of primary HNSCC cells grown in exosome-depleted FBS as described previously (21). Thirty ml of media was sequentially centrifuged at 300 × g for 10 minutes, 2,000 × g for 20 minutes, and 10,000 × g for 30 minutes at 4°C to remove cells and debris. Supernatants were ultracentrifuged at 100,000 × g for 70 minutes at 4°C using a Optima LE-80K ultracentrifuge (Beckman-Coulter, Brea, CA). Pellets were resuspended in 200 μl of sterile filtered PBS with a 0.22 μm syringe filter, washed by a second ultracentrifugation step, and stored at –80°C until further analysis (21).

#### Nanoparticle tracking analysis (NTA)

Exosome size and concentration were determined using a NanoSight NS300 system (Malvern) as previously described (21). Samples were diluted 1:100 to 1:1000 in 0.22 μm-filtered PBS to achieve 15–50 particles per frame. Each sample was analyzed five times for 30 seconds, and data were processed using NTA software.

#### Transmission electron microscopy

Morphological characterization of exosomes was performed by negative staining transmission electron microscopy (TEM) as previously described (21). Exosome suspensions (3 μl) were applied to glow-discharged carbon-coated grids (Electron Microscopy Sciences, Hatfield, PA), washed with 2% uranyl acetate, and imaged on a Talos L120C 120 kV TEM equipped with a Ceta 16M CMOS detector (ThermoFisher, Waltham, MA). Images were acquired with 1-second exposure times and analyzed using ImageJ (NIH).

#### Protein quantification

Total exosomal protein was quantified using the Qubit Protein Assay (ThermoFisher) following exosome lysis in cold RIPA buffer as per manufacturer’s instructions (21).

### Multiplex ELISA

Exosome surface proteins were quantified in multiplex using the U-PLEX assay platform (Meso Scale Discovery, MSD, Rockville, MD, USA), which enables the simultaneous detection of multiple analytes in a single well via electrochemiluminescence (ECL)(22). In this study, custom multiplexed capture antibody panels (Table S1) were used to detect immune checkpoint proteins, PD1 ligands (PDL1 and PDL2), TIGIT ligands (CD112, CD113, and CD155), and adenosine-generating enzymes (CD73 and CD39), and tetraspanins CD63, CD9, and CD81 on the surface of intact exosomes. Commercial antibodies against PDL2, CD112, CD113, and CD155 (Table S1) were biotinylated using the EZ-Link™ Sulfo-NHS-Biotin reagent (ThermoFisher) following the manufacturer’s protocol. Biotinylated R-PLEX antibodies for PDL1, CD63, CD9, and CD81 were obtained directly from MSD. U-PLEX linkers were first coupled to biotinylated capture antibodies to generate unique capture complexes, which were then added to streptavidin-coated 96-well U-PLEX plates, where they self-assembled onto predefined array elements. For each assay, intact exosomes corresponding to 20 μg total protein were used. Exosomes isolated from HNC208, HNC285, and HNC365 cells were added to the wells and incubated overnight (for immune checkpoint detection) or for 1 h (for tetraspanin detection). Control wells containing only the exosome resuspension medium were included to assess the background signal. After a blocking step, standards and samples were incubated to allow target analytes to bind their respective capture antibodies. Following washing to remove unbound material, a SULFO-TAG–labeled detection antibody cocktail specific for CD63, CD9, and CD81 (MSD) was added. After the final wash, MSD Read Buffer was added, and the plates were read using the MESO QuickPlex SQ 120MM instrument. Data were analyzed using MSD Discovery Workbench software. ECL signals exceeding the mean plus two standard deviations of the background (representing a 95% confidence interval) were considered indicative of positive analyte detection.

### PBMC isolation

Peripheral blood mononuclear cells (PBMCs) were isolated from discarded blood units from healthy individuals (healthy donors, HD) who donated blood at the Hoxworth Blood Center (University of Cincinnati) by Ficoll-Paque density gradient centrifugation (Cytiva Life Sciences, Marlborough, MA, USA) (19). Demographic information for these donors was not available.

### CD8 isolation and activation

CD8^+^ T cells were isolated from PBMCs by negative selection using the EasySep Human CD8^+^ T Cell Enrichment Kit (STEMCELL Technologies Inc.) (19). The CD8^+^ T cells were maintained in RPMI 1640 medium supplemented with 10% human serum, penicillin (200 U/ml), streptomycin (200 μg/ml), 1 mM L-glutamine, and 10 mM Hepes. For all experiments, cells were activated in a cell culture dish pre-coated with mouse anti-human CD3 antibody (10 μg/ml) (BioLegend, RRID: AB_11150592) and mouse anti-human CD28 antibody (10 μg/ml) (BioLegend, RRID: AB_11148949) for 72 to 96 h at 37°C, except where specified.

### Nanostring

Transcriptomic profiling of activated PBMCs (∼1 × 10⁶ cells/condition) from healthy donors (HDs) in the absence or presence of exosomes (1.04 x 10^9^ particles/ ml) was conducted using the NanoString nCounter® system (NanoString Technologies) (23). Total RNA was extracted with the E.Z.N.A. Total RNA Isolation Kit (Omega Bio-tek) and quantified by NanoDrop Lite plus spectrophotometer (ThermoFisher Scientific). RNA integrity was assessed using an Agilent 2100 Bioanalyzer (Agilent Technologies) and concentrated with the RNA Clean & Concentrator Kit (Zymo Research). Samples met quality thresholds of 260/280 ratio > 1.8, 260/230 ratio > 2.0, and DV200 > 30%.

One hundred nanograms of RNA per sample were hybridized to the PanCancer Immune Profiling CodeSet (NanoString), which targets 773 immune-related genes and 12 housekeeping genes. Hybridization and scanning were performed according to the manufacturer’s instructions using the NanoString nCounter® FLEX system. Each sample was processed in a multiplexed reaction alongside reference probes, including positive and negative controls and housekeeping genes, for normalization.

#### Nanostring Data Analysis

Raw gene expression data (counts) were calculated using the Basic Analysis feature of nSolver software (version 4.0, NanoString). The counts were normalized to the geometric means of internal housekeeping gene probes and subsequently imported into the nSolver Advanced Analysis plug-in (version 2.0, NanoString) for cell type profiling. This approach quantifies cell populations by employing marker genes as reference genes (Table S2), which are expressed stably and specifically in certain cell types. The scores for the abundance of immune cell types were determined using the log2 scale expression of their specific genes (22). Using the normalized gene expression counts, differentially expressed genes (DEGs) were identified with the ‘DESeq2’ package from R version 4.0.3 (24,25). Genes were considered statistically significant at log_2_fold change of 1 and -1, and a (non-adjusted) p-value ≤ 0.05 (-log_10_pvalue = 1.3). These genes were visualized through the ‘EnhancedVolcano’ package from R version 4.0.3 (25). Significant DEGs were subjected to pathways and biological processe enrichment analysis using piNET and gene ontology. The DEGs were further analyzed as per the workflow in (26), for gene-gene interactions using STRING-DB version 12 (https://string-db.org/) (accessed on February 14, 2025) and annotated according to the pathways they were enriched in, and visualized using Cytoscape v3.9.1.

### RT-qPCR

Total RNA was extracted from activated healthy donor PBMCs (∼1 × 10⁶ cells/condition) in the absence or presence of exosomes (1.04 x 10^9^ particles/ ml) using the E.Z.N.A. Total RNA Kit (Omega Bio-tek). 300 ng of RNA was transcribed into cDNA employing the LunaScript RT SuperMix Kit (NEB), following the manufacturer’s instructions. RT-qPCR was conducted utilizing TaqMan Gene Expression Assays (ThermoFisher) to quantify *KCNA3* (Hs00704943_s1), *KCNN4* (Hs01069779_m1), *ORAI1* (Hs03046013_m1), *STIM1* (Hs00963373_m1), *CALM1* (Hs00300085_s1), *CALM2* (Hs00830212_s1), *CALM3* (Hs00968732_g1), with 1*8S rRNA* (Hs99999901_s1) serving as the reference control. Each reaction comprised 30 ng of cDNA, 1X TaqMan Master Mix, and 1 μL of each primer probe, performed in triplicate on a QuantStudio Real-Time PCR System. C_T_ values were analyzed using QuantStudio software, with gene expression normalized to 18S rRNA via the ΔΔCT method. Relative expression (2^−ΔΔC^_T_) was calculated in comparison to healthy donor PBMCs, as previously described (23).

### Flow Cytometric Phenotyping of PBMCs

Cryopreserved PBMCs (∼1 × 10⁶ cells/condition) from HDs were thawed and rested overnight in T cell medium. The next day, the cells were activated for 72 h with plate bound anti-CD3 and anti-CD28 antibodies in the absence or presence of exosomes (1.04 x 10^9^ particles/ ml). The cells were then resuspended in 100 μl PBS, and stained with Zombie UV Live/Dead dye (BioLegend) as per the manufacturer’s instructions, washed with staining buffer (BioLegend), fixed in 4% paraformaldehyde for 30 minutes and stained overnight at 4°C with guinea pig anti-Kv1.3 and mouse anti-KCa3.1 (ATTO-488) primary antibodies (Alomone Labs) in the dark. The following day, cells were washed and incubated with Alexa Fluor 555-conjugated goat anti-guinea pig IgG secondary antibody (Thermo Fisher), followed by surface staining with an antibody cocktail targeting CD69, CD3, CD4, CD8, CD19, CD56, CD16, and CD14. A full list of antibodies is provided in Table S3. The specificity of Kv1.3 antibodies used in this study was validated previously (27). Samples were acquired on a BD LSRFortessa flow cytometer and analyzed using FlowJo software (BD Biosciences). Unstained cells, single-stained controls, and UltraComp eBeads (ThermoFisher) were used for compensation.

### Intracellular Ca²⁺ Measurement

Intracellular calcium (Ca²⁺) flux was assessed using the Ca²⁺ add-back method as previously described (27). Briefly, 1 × 10⁶ freshly isolated CD8⁺ T cells, treated with exosomes (1.04 × 10⁹ particles/ml) or PBS (control), and activated with anti-CD3/CD28 antibodies for 72 hours, were loaded with the ratiometric Ca²⁺ indicator Indo-1 AM (2 mg/ml, 1:1000 dilution; ThermoFisher) in the presence of 0.015% Pluronic F-127 (ThermoFisher). Cells were incubated in Hank’s Balanced Salt Solution (HBSS) supplemented with 1 mM CaCl₂, 1 mM MgCl₂, and 1% FBS for 30 minutes at 37°C. Following dye loading, cells were washed three times with HBSS containing 10 mM HEPES (pH 7.0) and 1% FBS. Prior to acquisition, cells were resuspended in Ca²⁺-free HBSS/HEPES buffer containing 0.5 mM EGTA (pH 7.4) and maintained at 37°C. Real-time Indo-1 fluorescence ratios (violet/blue emission) were acquired using a BD LSRFortessa flow cytometer with previously optimized instrument settings (23). To assess Ca²⁺ influx, cells were first stimulated with thapsigargin (1 μM, MilliporeSigma) in Ca²⁺-free buffer to deplete intracellular Ca²⁺ stores and thereby activate CRAC channels. This was followed by the addition of 2 mM extracellular Ca²⁺, allowing Ca²⁺ to enter the cells through the open CRAC channels. Data were analyzed using FlowJo software (BD Biosciences), and intracellular Ca²⁺ flux was quantified as the fold change in Indo-1 fluorescence ratio, calculated by dividing the post–Ca²⁺ addition peak by the baseline fluorescence in Ca²⁺-free conditions.

### Electrophysiology

K^+^ currents were recorded in whole-cell patch configuration. The external solution was (in mM): 140 NaCl, 4.5 KCl, 2 CaCl_2_, 1 MgCl_2_, 10 Hepes, pH 7.4. The pipette solution was (in mM): 145 K aspartate, 8.5 CaCl_2_, 10 EGTA, 2 MgCl_2_ and 10 Hepes, pH 7.2, with an estimated free Ca^2+^ concentration of 1 μM (19). K^+^ current was measured in voltage-clamp mode and induced by ramp depolarization from −120 mV to +40 mV, 200 ms duration, every 10 s, −80 mV holding potential (HP). KCa3.1 channel activity was quantified between −100 mV and −80 mV by calculating the ratio of the linear portion of the macroscopic current slope to the slope of the ramp voltage stimulus, after subtraction of the leak current (19). Kv1.3 channel activity was assessed at +40 mV using the same ramp protocol after subtracting the KCa3.1 current estimated through linear regression. The digitized signals were stored and analyzed using pClamp 9 software (Axon Instruments). Data were corrected for a junctional potential of -10 mV (28).

### LEGENDplex™ Cytokine and Cytotoxicity Assay

Cytokine and cytotoxic marker secretion by CD8^+^ T cells was assessed using the LEGENDplex™ Human CD8/NK Panel (BioLegend; Cat# 741187), which quantifies IL-17A, IL-2, IL-4, IL-10, IL-6, TNF-α, soluble Fas (sFas), soluble Fas-ligand (sFasL), IFN-γ, granzyme A, granzyme B, perforin, and granulysin. The assay was performed according to the manufacturer’s instructions. Briefly, 1 × 10⁶ CD8^+^ T cells from HDs were activated with anti-CD3/CD28 antibodies for 72 h, in the presence or absence of exosomes (0.5 x 10^9^ particles/ml), respectively. Supernatants (25 μl) were collected and the assay was performed in duplicate in a 96-well format according to the manufacturer’s instructions. Samples were acquired on a Cytek® Aurora spectral flow cytometer equipped with a yellow-green laser and an automated 96-well plate reader, using SpectroFlo® software and manufacturer-recommended settings. Data were analyzed using LEGENDplex™ Data Analysis Software. Results were reported as a fold change in mean fluorescence intensity (MFI) relative to vehicle-treated controls.

### IFN-γ determination

Cryopreserved PBMCs were thawed and rested overnight in T cell medium. CD8^+^ T cells were isolated and 1 × 10⁶/ml cells activated for 72 h with anti-CD8/CD28 antibody in the absence or presence of exosomes (0.13 × 10⁹ particles/ml) +/− 1 μM NS309 (MilliporeSigma). IFN-γ levels in the culture supernatants were measured using a Human IFN-γ Uncoated ELISA Kit (Thermo Fisher, Catalog # 88-7316-88) as per the manufacturer’s protocol.

### Statistical Analysis

Statistical analyses were performed using Student’s t-test (paired or unpaired). The normality of sample distribution was assessed by the Shapiro–Wilk test, and where the samples failed normality, comparisons were performed by the Mann–Whitney rank sum test. Statistical analysis was performed using GraphPad Prism 9.0 (GraphPad Software LLC, Boston, MA, USA). P ≤ 0.05 was defined as statistically significant. Appropriate statistical tests and corresponding values are described in individual figure legends.

### Data availability

All data needed to evaluate the conclusions in the paper are presented in the paper and/or Supplementary Materials. Additional data related to this study may be requested from the authors.

## Results and Discussion

### Exosomes from patient-derived HNSCC cells carry immune checkpoint inhibitor proteins on their surface

In this study, we tested the immune effects of exosomes isolated from patient-derived HNSCC cells and investigated the mechanisms mediating these effects. HNSCC primary cell cultures were obtained from resected HPV-negative (p16-negative) and PDL1-positive tumors from three treatment naïve patients (HNC208, HNC285 and HNC365) (21). Exosomes were isolated from the conditioned media by ultracentrifugation and characterized in accordance with the requirements described by the International Society for Extracellular Vesicles (ISEV) (29,30). We measured the size and integrity of the exosomes by NTA (Fig1A) and TEM (Fig1B), with results consistently showing them to be small extracellular vesicles (exosomes) exhibiting diameters less than 200 nm. The presence of tetraspanins (CD9, CD63, and CD81) on the surface of HNSCC exosomes was detected by multiplex ELISA (Fig1C). This method of tetraspanin detection was validated with on-bead flow cytometry (FigS1A) (21).

To determine whether known immunosuppressive molecules of the TME were present on exosomes, we measured the levels of PD1 ligands (PDL1 and PDL2), TIGIT ligands (CD155, CD112, CD113) and adenosine generating enzymes (CD39 and CD73) on intact exosomes from HNC208, HNC285, and HNC365 cells by multiplex ELISA. While all exosomes carried PD-L1 on their surfaces, as expected given their derivation from PD-L1⁺ tumors (21), their overall composition varied between patients (Fig1D). HNC208 showed detectable levels of all immune inhibitory proteins tested, whereas the profiles for HNC285 and HNC365 exosomes were different, with HNC365 exosomes showing a signature enriched in CD73. To verify the multiplex ELISA method, we utilized a PDL1 knockout Cal27 cell line we generated by the CRISPR/Cas9 system and confirmed the detection of PDL1 in wild-type, but not in knockout, cells by both multiplex ELISA and on-bead flow cytometry (FigS1B). Given the immunosuppressive potential of HNSCC-derived exosomes, we proceeded to investigate their effects on human immune cells.

### HNSCC-derived exosomes inhibit the anti-tumor capabilities of cytotoxic T cells

To test the immunosuppressive effects of tumor-derived exosomes, we treated PBMCs from HDs with HNSCC exosomes (1.04 x 10^9^ exosomes/ml), followed by activation with anti-CD3/CD28 antibodies. This exosome concentration was chosen to corresponds to the plasma exosome concentration in early stage HNSCC patients (1.05 x 10^9^ particles/ml) (31) and used in all experiments unless otherwise stated. Transcriptomic analysis using the NanoString platform with the Pan Cancer Immune Profiling panel revealed that exosomes reduced the abundance of anti-tumor cells like cytotoxic cells, in particular CD8^+^ T cells and CD56^dim^ NK cells (the highly cytotoxic NK cell subset found in tumors), Th1 CD4^+^ T cells and dendritic cells, and pro-tumor regulatory T (Treg) cells and mixed-function macrophages (this method did not allow us to distinguish between M1 and M2 macrophages) (Fig2A; FigS2) (2). HNSCC exosomes increased mast cells (mixed tumor function) and B cells (anti-tumor). Differential gene expression analysis, GSEA, and STRING showed that exosomes inhibited leukocyte functions, including immune responses associated with cytotoxicity, chemokines, and NK and T cell functions (Fig2B, Table S4). Specifically, exosomes downregulated genes involved in antigen presentation, processing and cell adhesion (*HLA-DQA1*, *HLA-DQB1*, and *CD276*), cytokine and chemokine-related signaling (*IL-9*, *IL-6*, *IL-11*, *IL-23A*, and *TLR8*), and tumor necrosis factor (*TNF*) signaling (Fig2C-D). Importantly, HNSCC exosomes downregulated the Ca^2+^-dependent NF-kB signaling pathway, critical for cytotoxic CD8^+^ T cell function and effector cytokine production (e.g., IFN-γ) (13,32).

Further experiments conducted using a flow cytometry-based multiplex immunoassay, showed that exosomes (0.5 x 10^9^ particles/ml) inhibited the release of effector cytokines (IL-2, TNF-α, IFN-γ), Fas ligand (FasL) and cytotoxic molecules (granzyme B, perforin, and granulysin) from activated human CD8⁺ T cells (Fig2E; FigS3). This exosome concentration falls within the range of maximal effect, based on previously generated dose-response curves assessing the impact of HNSCC-derived exosomes from these same patients on IFN-γ production by healthy donor PBMCs (21). These effector cytokines and cytotoxic molecules altered by HNSCC exosomes are necessary for effective CD8^+^ T cell tumor killing. Fas-FasL interaction also contributes to effective antitumor effects as it mediates off-target bystander killing of antigen-negative tumor cells (33). Overall, these studies demonstrate that HNSCC exosomes significantly impair immune cell function and provide a central role for NF-kB signaling pathway in mediating these effects. NF-kB signaling, along with the expression of IL-2, TNF-α, IFN-γ, and FasL are regulated by Ca^2+^ (13).

Ca^2+^ influx is one of the earliest events in TCR stimulation and serves as a precise regulator of gene expression. The amplitude and frequency of Ca^2+^ oscillations can selectively activate transcription factors such as NF-κB and NF-AT (13,32). NF-κB is particularly responsive to low-frequency Ca²⁺ oscillations because its nuclear translocation is regulated by the slow degradation and resynthesis of its inhibitor, IκB. Upon intracellular Ca²⁺ elevation and phorbol ester stimulation, IκB is phosphorylated and degraded, enabling NF-κB to enter the nucleus. There, it remains active until newly synthesized IκB accumulates and binds NF-κB, sequestering it back in the cytoplasm. Because IκB resynthesis is a slow process—taking tens of minutes—the nuclear exit of NF-κB is similarly delayed. This explains the sustained NF-κB activity in response to infrequent Ca²⁺ spikes (32).

In addition to transcriptional regulation, Ca²⁺ is also essential for the exocytosis of cytotoxic granules containing perforin and granzymes (13). When Ca²⁺ enters the cell upon immune synapse formation, it activates signaling pathways that lead to the fusion of granules with the plasma membrane, enabling their release to kill target cells. Despite its central role in T cell effector functions, the impact of exosomes on Ca²⁺ signaling remains poorly understood. Ca^2+^ fluxes in T cells are governed by the function of ion channels that act in concert to regulate the membrane potential necessary to establish the appropriate driving force for Ca^2+^ influx through Ca^2+^ channels (13).

### Inhibition of KCa3.1 channels mediates the effects of HNSCC exosomes on CD8^+^ T cells Ca^2+^ fluxing abilities and downstream effector functions

We conducted RT-qPCR in PBMCs to measure the effect of HNSCC-exosomes on the gene expression of the canonical ion channels regulating Ca^2+^ signaling in activated T cells: KCa3.1 (encoded by the *KCNN4* gene), Kv1.3 (*KCNA3* gene), and Orai1 and Stim1 (the subunits forming CRAC channels and encoded by *ORAI1* and *STIM1*). Exosomes significantly increased and decreased *KCNN4* and *KCNA3*, respectively, while had no effect on *ORAI1* and *STIM1* (Fig3A). Flow cytometry experiments of human PBMCs activated with anti-CD3/CD28 antibodies with and without HNSCC exosomes revealed no alterations in either K^+^ channel protein levels or CD69 (activation marker) in CD3^+^, CD4^+^, CD8^+^ and NK cells apart from a decrease in Kv1.3 protein levels in CD8^+^ T cells (Fig3B-E and FigS4-S5). The gating strategy of these experiments is shown in FigS4. Exosome exposure did not alter the cell viability (FigS6A) or the percentage of CD3^+^, CD4^+^, CD8^+^, or cytotoxic NK (CD56^dim^CD16^+^) immune cell subsets (Fig. S6B). However, electrophysiological experiments showed that HNSCC exosomes selectively inhibited KCa3.1, but not Kv1.3 channel activity in CD8^+^ T cells (Fig4A).

The activity of KCa3.1, a Ca^2+^ activated K^+^ channel that opens in response to an increase in intracellular Ca^2+^ triggered by TCR stimulation, is regulated by calmodulin, a Ca^2+^-sensor constitutively bound to the C-terminal domain of the channel, and necessary to confer the channel’s Ca^2+^-sensitivity (34). Calmodulin is encoded by 3 different genes (*Calm1*, *Calm2* and *Calm3*) (35). *Calm2* and *Calm3* genes were downregulated by exosomes leading to a 30% reduction in calmodulin protein levels (Fig4B-C). We have reported that, while KCa3.1 surface protein levels are the same in healthy donor and HNSCC patients’ circulating T cells, there is areduction in KCa3.1 activity and membrane calmodulin in HNSCC that leaves a high proportion of KCa3.1 channels devoid of calmodulin and, therefore, not functional (19,35). The same phenotype is reproduced in healthy peripheral T cells by HNSCC exosomes, suggesting that exosomes may be responsible for the KCa3.1 channel defects of circulating T cells in HNSCC patients that limits the ability of these cells to infiltrate the TME.

The decrease in functionally competent KCa3.1 channels induced by exosomes in CD8^+^ T cells is associated with a reduction of Ca^2+^ fluxing ability in these cells (Fig4D). Flow cytometry studies with the Ca^2+^ sensitive dye Indo-1 showed that HNSCC exosomes significantly suppress the Ca^2+^ fluxing abilities of CD8^+^ T cells by 49%. Ca^2+^ fluxes were elicited by thapsigargin (TG)/Ca^2+^ add-on method (27). This method allows measuring changes in intracellular Ca^2+^ levels that depend exclusively on the ionic events downstream to the T cell receptor. Further IFN-γ measurement studies of CD8^+^ T cells exposed to 0.13 x10^9^ exosomes/ml, showed that the KCa3.1 activator NS309 restores the ability of CD8^+^ T cells to secrete IFN-γ in the presence of HNSCC exosomes (Fig. 4E). This concentration of exosomes was used because it corresponds to the IC₅₀ (half maximal inhibitory concentration) of exosomes for IFN-γ release, as previously determined in our ELISA-based assays (21).

These data provide evidence of the key role of KCa3.1 channels in the immune suppressive effects of HNSCC exosomes on CD8^+^ T cells. KCa3.1 channels are also inhibited by other immunosuppressive factors present in the TME including adenosine, PGE2 and PDL1 (36). We show here that HNSCC exosomes carry adenosine producing enzymes known to be functional in exosomes from other types of cancers, i.e. capable of producing adenosine from ATP (37). Adenosine inhibits KCa3.1 via PKA (36). They also carry PDL1 and TIGIT ligands albeit with distinct patient-to-patient variability (Fig1D). PDL1 inhibits KCa3.1 channels in T cells via PI3K (36). TIGIT has been shown to signal through PI3K (38), suggesting a common mechanism of KCa3.1 modulation for PD1 and TIGIT ligands. Given the presence of these proteins on the surface of exosomes, it is likely that exosomes engage these signaling pathways to suppress KCa3.1 activity in cytotoxic T cells. Overall, to the best of our knowledge, this is the first study to show that the immunosuppressive effects of multiple elements of the TME converge on KCa3.1 channels, positioning these channels as strategic targets to enhance immune surveillance in cancer.

**Fig. 1:**
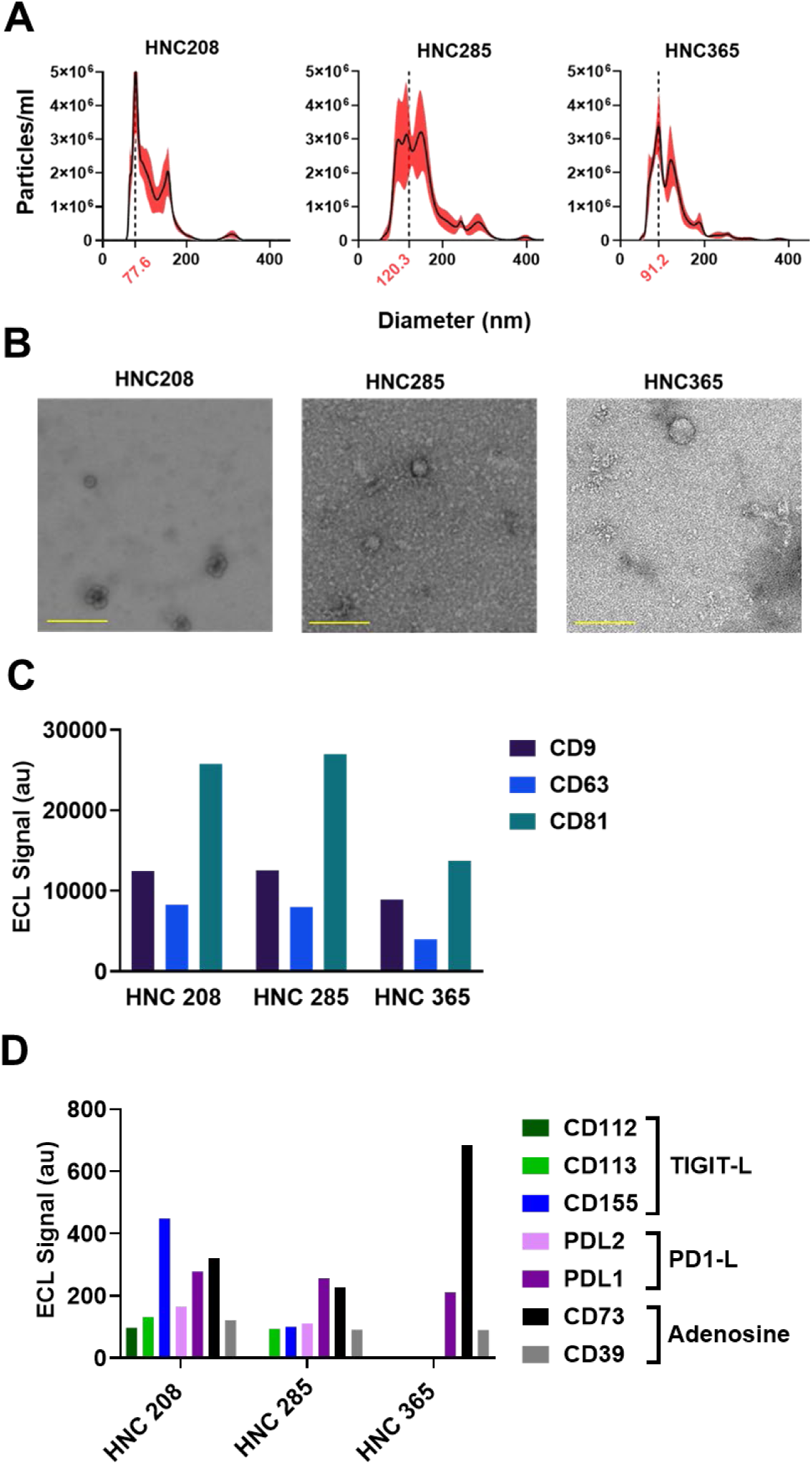
Characterization of exosomes derived from HNSCC cells. (A) Size distribution and concentration of exosomes isolated from the supernatant from three HNSCC patient derived cancer cells measured by NTA. Values are represented as mean ± SEM from 5 independent captures. (B) Representative transmission electron micrographs showing exosomes isolated from cancer cell culture supernatants from HNSCC patients (scale bar = 200 nm). (C) Tetraspanins CD63, CD81, and CD9 were detected on the surface of intact exosomes from three HNSCC patients by multiplexed ELISA. Exosomes were captured by antibodies to single tetraspanins and detected by a signal from a cocktail of anti-tetraspanin antibodies (CD63, CD81, and CD9). Each sample was run in duplicate on the same plate. (D) Surface protein phenotyping of intact exosomes by multiplexed ELISA showing the abundance of PD1 ligands (PD1-L, PDL1 and PDL2), TIGIT ligands (TIGIT-L, CD112, CD113, CD155) and adenosine generating enzymes (CD73 and CD39) on exosomes released by different patient cancer cells. Exosomes were captured by antibodies against immune checkpoint proteins and adenosine generating enzymes and detected by a cocktail of anti-tetraspanin antibodies (CD63, CD81, and CD9). Each sample was run in duplicate on the same plate.

**Fig 2:**
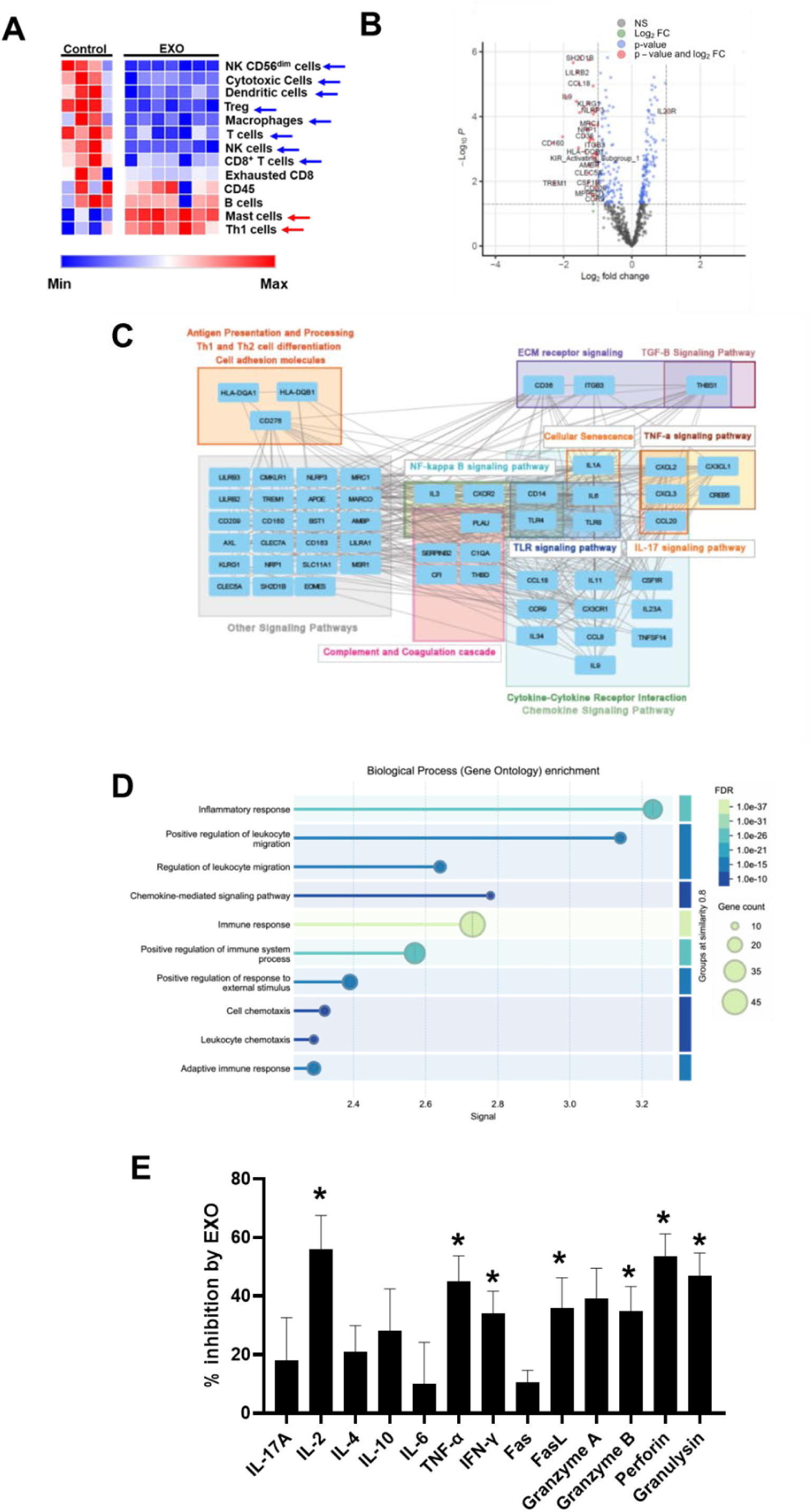
Transcriptomic and secretory profiles of immune cells treated with HNSCC exosomes. (A) Heatmaps generated from NanoString transcriptomic data depicting the relative immune cell abundance in PBMCs from untreated healthy donors (n=4; Control) and from healthy donors treated with exosomes (n=7; EXO) at a concentration of 1.04×10⁹ particles/ml. The EXO group included PBMCs from four healthy donors treated with exosomes derived from three different HNSCC patients (HNC208, HNC285, and HNC365); three donors were each treated with exosomes from two different patients. Significance between groups was determined by either Mann-Whitney rank sum test (cytotoxic cells, B cells, and CD8 T cells), or unpaired t-test and is shown by arrows. Blue and red arrows indicate decrease and increase in cell abundance, respectively. Bar graphs comparing the immune cell abundance between the two groups along with the numerical values for statistical significance are shown in Fig. S2. (B) Volcano plot showing Differential Gene Expression analysis of NanoString Pan Cancer Immune Profiling data between exosome treated PBMCs and untreated PBMCs (Control). When healthy donor PBMCs were treated with exosomes, 59 genes were downregulated, and 4 genes were upregulated of 770 immune-related genes, at 2-fold change and p<0.05.The differentially expressed genes are listed in Table S4. (C) Gene-Gene Interaction Network of significant differentially expressed genes (Table S4). (D) Significant differentially expressed genes were annotated for their gene ontology. The downregulated genes are involved in regulating inflammatory response, leukocyte migration, NK and cytotoxic T cell function, chemokine and cytokine signaling. (E) Multiplex cytokine release assay showing the percentage inhibition of individual cytokine levels in activated CD8⁺ T cells treated with exosomes (0.5 ×10⁹ particles/ml) from HNC208, HNC285, and HNC365, compared to untreated controls. Data were obtained from six healthy donors (n=6), with the same donors used for both treated and control conditions. * denotes significance comparing control and treatment groups, and error bars represent SD. Specific p-values (calculated between the groups using the Mann-Whitney rank sum test) are reported in Fig S3.

**Fig. 3:**
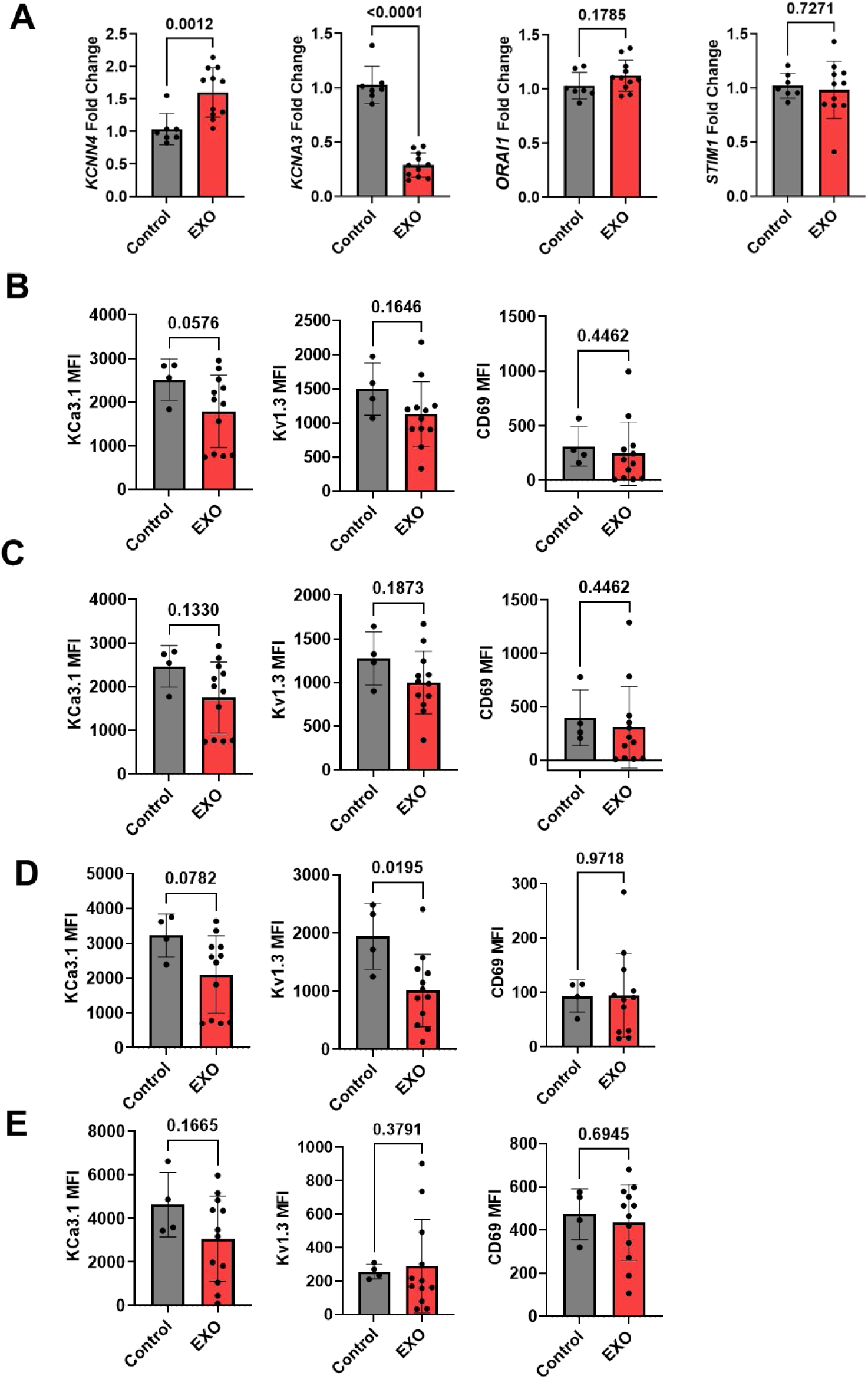
Alterations in ion channel expression in immune cells treated with HNSCC exosomes. (A) Fold change in mRNA abundance of KCa3.1 *(KCNN4*), Kv1.3 (*KCNA3*), Orai (*ORAI1*) and Stim1 (*STIM1*) in PBMCs from healthy donors (n=7) treated with HNC208, HNC285 and HNC365 exosomes (EXO, 1.04 X 10^9^ particles/ ml) was determined by RT-qPCR. PBMCs from some donors in the EXO group were each treated with exosomes from two different HNSCC patients. Each sample was run in triplicate. *18S rRNA* was used as the housekeeping gene. Data were normalized to untreated PBMCs (Control). Bars represent means ± SD, and symbols represent individual treatment conditions. Significance was determined by unpaired t-test (*ORAI1* and *STIM1*) and Mann-Whitney rank sum test (*KCNN4* and *KCNA3*). (B-E) Mean fluorescence intensity (MFI) of KCa3.1, Kv1.3, and CD69 in CD3^+^ (B), CD4^+^ (C), CD8^+^ (D), and CD56^dim^CD16^+^ NK cells (E) from four healthy donors treated with HNSCC exosomes from HNC208, HNC285, and HNC365 (same protocol as in A, some donors were treated with exosomes from multiple HNSCC patients). Untreated PBMC were used as controls. Significance was determined by unpaired t-test and Mann-Whitney rank sum test (CD3^+^CD69, CD4^+^KCa3.1, CD4^+^CD69, NK^+^Kv1.3, NK^+^CD69). Bars represent means ± SD, symbols represent individual treatment conditions.

**Fig. 4:**
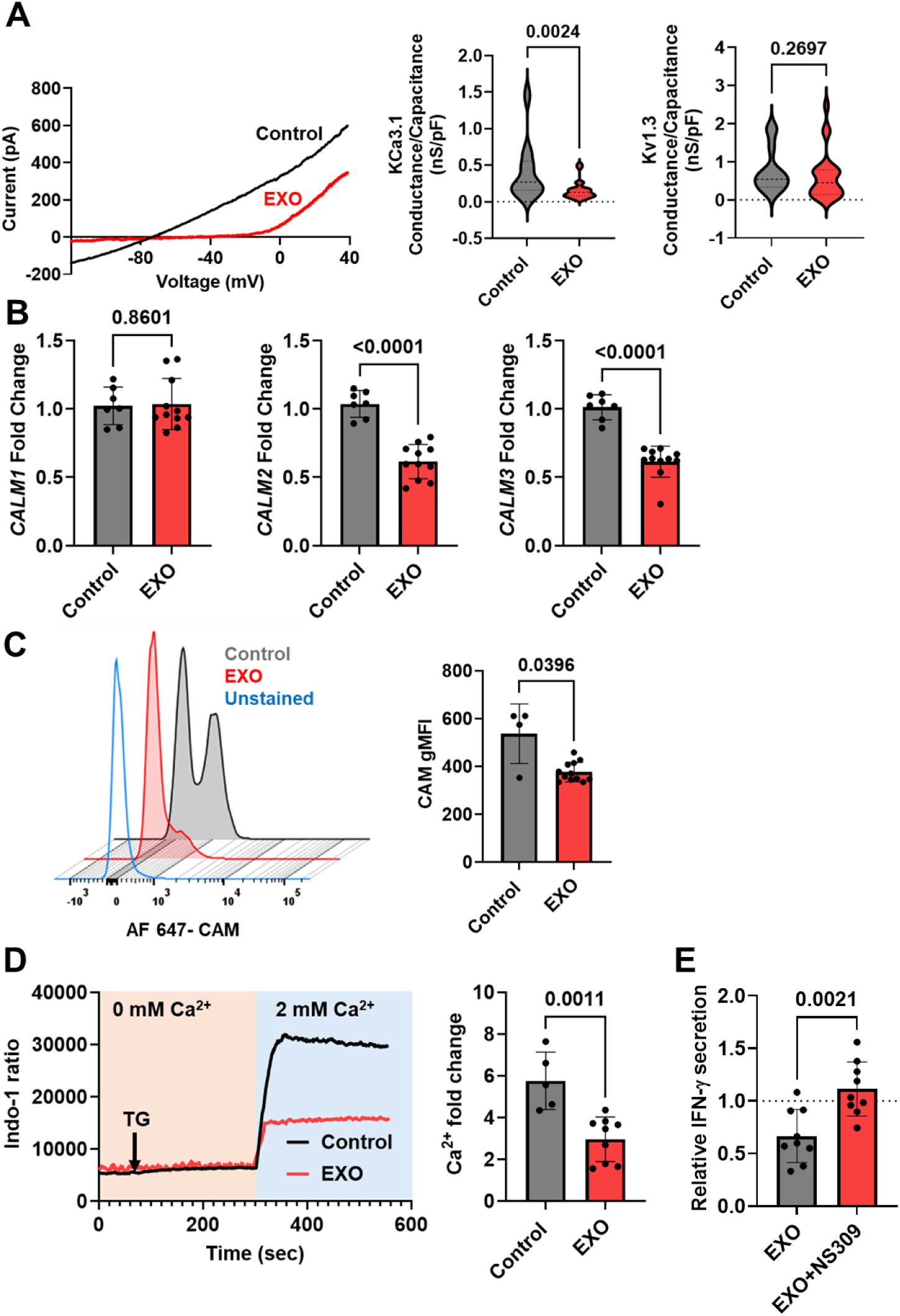
Inhibition of KCa3.1 channel activity and Ca^2+^ fluxing abilities of CD8^+^ T cells by HNSCC exosomes. (A) (Left) Representative KCa3.1 currents recorded in whole-cell voltage-clamp configuration in activated CD8^+^ T cells from a healthy donor in the absence (Control) or presence of HNSCC exosomes (EXO; 1.04 X 10^9^ particles/ ml). (Right) Violin plots of KCa3.1 and Kv1.3 conductance (normalized to cell capacitance) in activated CD8^+^ T cells from five healthy donors in the absence (Control) or presence of HNSCC exosomes (EXO) from HNC208, HNC285, and HNC365 (n= 21 cells for control and 23 cells for EXO; some donors’ T cells were treated with exosomes from multiple HNSCC patients). Significance was determined using the Mann-Whitney rank sum test. (B) Fold change in mRNA abundance of *CALM1*, *CALM2,* and *CALM3* in PBMCs from healthy donors (n=7) treated with HNC208, HNC285, and HNC365 exosomes (EXO; 1.04 X 10^9^ particles/ ml) was determined by RT-qPCR. The PBMCs from some donors in the EXO group were each treated with exosomes from two different HNSCC patients. Each sample was run in triplicate. *18S rRNA* was used as the housekeeping gene. Data were normalized to untreated PBMCs (Control). Bars represent means ± SD, and the symbols represent individual treatment conditions. Significance was determined using an unpaired t-test (*CALM2*) and Mann-Whitney rank sum test (*CALM1* and *CALM3*). (C) Representative flow cytometry staggered histograms showing CaM expression in CD8^+^ T cells from one healthy donor in the absence (Control) or presence of HNC 208, HNC 285, and HNC 365 exosomes (EXO; 1.04 X 10^9^ particles/ ml). The bar graph on the right shows the geometric mean fluorescence intensity (gMFI) for CaM expression in control and EXO-treated cells from four healthy donors. Bars represent mean ± SD; symbols represent individual treatment conditions (some donors were treated with exosomes from multiple HNSCC patients). Significance was determined by Mann-Whitney rank sum test. (D) Representative Ca^2+^ response (shown as a ratio of Indo-1 fluorescence at 400 and 480 nm) recorded in activated healthy donor CD8^+^ T cells in the absence (Control) or presence of HNC208, HNC285, and HNC365 exosomes (EXO; 1.04 X 10^9^ particles/ ml). Cells were loaded with Indo-1; and analyzed by flow cytometry, thapsigargin (arrow) was added in 0 mM Ca^2+^ to induce intracellular Ca^2+^ store depletion, followed by 2 mM Ca^2+^, which yielded a rapid Ca^2+^ influx (see Materials and Methods). Right panel shows the average peak Ca²⁺ fold change in CD8^+^ T cells from n = 5 donors. Bars represent mean ± SD; symbols represent individual treatment conditions (some donors were treated with exosomes from multiple HNSCC patients). Statistical significance was determined by unpaired t-test. (E) IFN-γ levels in healthy donor CD8^+^ T cells (n =3 healthy donors) treated with HNSCC exosomes (0.13 × 10^9^ particles/ml), isolated from supernatants of HNC208, HNC285, and HNC365 cells, and activated with plate-bound anti-CD3 and anti-CD28 antibodies for 72 h in the absence or presence of 1 μM NS309 (KCa3.1 activator). Each donor was exposed to HNC208, HNC285, and HNC365 exosomes, and the samples were run in duplicate on the same ELISA plate. Activated CD8^+^ T cells that were not exposed to exosomes were used as controls. For each experiment, IFN-γ levels in control samples were normalized to 1 (dotted line), and all other values were expressed relative to this baseline. Symbols represent individual treatment conditions; bars indicate mean ± SD. Significance was determined using a paired t-test.

## Supporting information

All supplementary data

## Acknowledgements

The authors would like to thank the patients and the healthy donors who participated in this study. We also thank the clinical coordinators at the University of Cincinnati Cancer Center’s Clinical Trials Office for helping collect patient samples. We appreciate the assistance of Ms. Damaris Kuhnell from the Department of Environmental & Public Health Sciences at the University of Cincinnati for her help with the nanoparticle tracking analysis. TEM was done using equipment from the Center for Advanced Structural Biology at the University of Cincinnati. We thank Dr. Desiree A Benefield, the Manager of the Center, for her help with the TEM experiments. We also thank Dr. Takahisa Nakamura (Division of Endocrinology at Cincinnati Children’s Hospital Medical Center), for providing NanoSight NS300. Multiplex ELISA assays (read on MESO QuickPlex SQ 120MM) and all flow cytometry experiments were performed using instruments maintained and operated by the Research Flow Cytometry Core, Division of Rheumatology, at Cincinnati Children’s Hospital Medical Center. We would like to thank Dr. Arpita Agrawal, Principal Field Application Scientist, Meso Scale Diagnostics LLC, for her technical assistance in optimizing the multiplex ELISA assays. RNA quality control was performed at Genomics, Epigenomics and Sequencing Core, Department of Environmental Health, University of Cincinnati. NanoString experiments were performed using equipment maintained by the Diagnostic Immunology Laboratory, Cincinnati Children’s Hospital Medical Center. PDL-1 knockout Cal27 cell lines (PDL1-KO) Cal 27 cells were generated with CRISPR-Cas9 system at Transgenic Animal and Genome Editing (TAGE) core at Cincinnati Children’s Hospital Medical Center. We are grateful to Dr. Roman Jandarov (Department of Biostatistics and Bioinformatics, University of Cincinnati) for his valuable counsel on statistical analysis. Paperpal, an AI-driven academic writing assistant, was used to enhance grammar and academic tone. No AI tools were used to generate scientific content, analyze data, or interpret the results. All scientific works were independently conceived, developed, and validated by the authors.

## Author Contributions

**Ameet A. Chimote**: Conceptualization, Study Design, Methodology, Investigation, Data Curation, Formal Analysis, Visualization, Supervision, Writing – Original Draft

**Abdulaziz O. Alshwimi**: Investigation, Formal Analysis

**Simran Venkatraman**: Formal Analysis, Visualization

**Jay Bhati**: Investigation

**Maria A. Lehn**: Investigation

**Somchai Chutipongtanate**: Study Design, Formal Analysis, Writing – Review & Editing

**Sarmistha Das**: Formal Analysis

**Susan Kasper**: Resources

**Shesh N. Rai**: Formal Analysis

**Scott M. Langevin**: Study Design, Writing – Review & Editing

**Trisha M. Wise-Draper**: Resources

**Laura Conforti**: Conceptualization, Supervision, Study Design, Formal Analysis, Funding Acquisition, Writing – Original Draft

**All the authors discussed the results and commented on the manuscript.**

