## Supplementary material for "Potassium channels mediate the inhibitory effect of exosomes on anti-tumor immunity in head and neck cancer": All supplementary data

**Supplementary Materials:**

**Supplementary Methods:**

**Cal27 cell culture**

Cal27 cells were obtained from American Tissue Culture Collection (ATCC, Manassas, VA, USA), grown as described previously in DMEM with 10% FBS, 1% penicillin–streptomycin, 1% minimum essential amino acids, 6 mM L-glutamine, and 1% sodium pyruvate in the presence of 5% CO2 at 37 °C in a humidified incubator (21). Mycoplasma testing was performed routinely using MycoStrip® mycoplasma detection kit (Invivogen, Catalog# rep-mys-50) and was negative for all cultures. PDL-1 knockout Cal 27 cell lines (PDL1-KO) were generated with CRISPR-Cas9 system at Transgenic Animal and Genome Editing (TAGE) core at CCHMC. Knockouts were validated by the core using RT-PCR. For the final two passages before exosome isolation, cells were cultured in media with 5% exosome-depleted FBS. Exosomes were isolated from the conditioned media of these cultured cells by ultracentrifugation (see Materials and Methods), and characterized as described previously (21).

**On-bead flow cytometry.**

Surface marker expression on exosomes was analyzed using on-bead flow cytometry as detailed previously (21). Exosomes (20 μg protein, measured with Qubit Assay, as described in Materials and Methods) were captured on anti-CD63 antibody-coated Dynabeads (Thermo Fisher) overnight at 4°C in isolation buffer (PBS + 0.1% BSA, 0.22 μm-filtered). Exosomes were stained with PE-conjugated antibodies against CD63 (RRID: AB_10896786), CD9 (RRID: AB_2075893), and CD81 (RRID: AB_10642024) (BioLegend). Controls included PE-conjugated IgG (RRID: AB_326435) and unstained beads. Data were acquired on a BD LSRFortessa cytometer and analyzed using FlowJo software (BD Biosciences).

**Supplementary Figures and Tables**

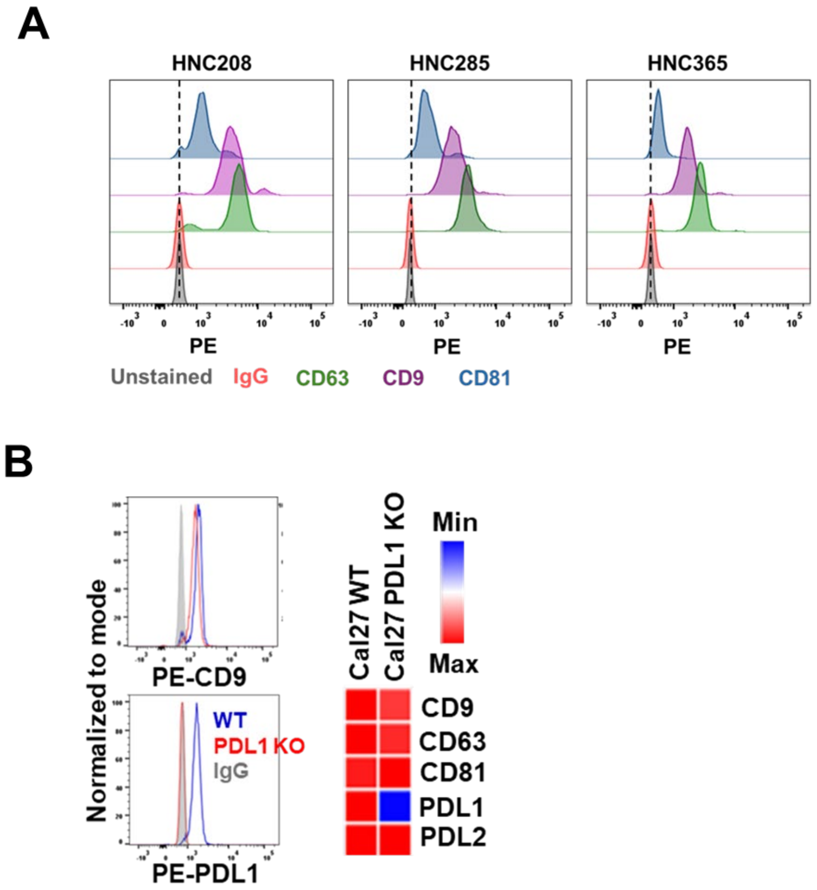


**Fig. S1 Validation of exosome characterization methodology** (A) Staggered histograms showing CD63, CD9, and CD81 abundance in exosomes isolated from HNSCC cell conditioned media that were captured on CD63-antibody-coated beads. Unstained and isotype control (IgG) stained beads were used as negative controls. (B) (Left) Overlay histograms showing CD9 (Top) and PDL1 (bottom) abundance in exosomes isolated from wild type (WT) and PDL1 knockout (PDL1-KO) Cal 27 cells that were captured on CD63-antibody-coated beads. Isotype control (IgG) stained beads were used as negative controls. (Right) Heatmap showing surface protein phenotyping of exosomes from WT and PDL1-KO Cal27 cells showing the abundance of tetraspanins (CD9, CD63 and CD81) and PD-1 ligands (PDL-1 and PDL-2) on intact exosome surface by multiplexed ELISA. Exosomes were captured by antibodies and detected by a signal from a cocktail of anti-tetraspanin antibodies (CD63, CD81, and CD9). Each sample was run in duplicate on the same plate.

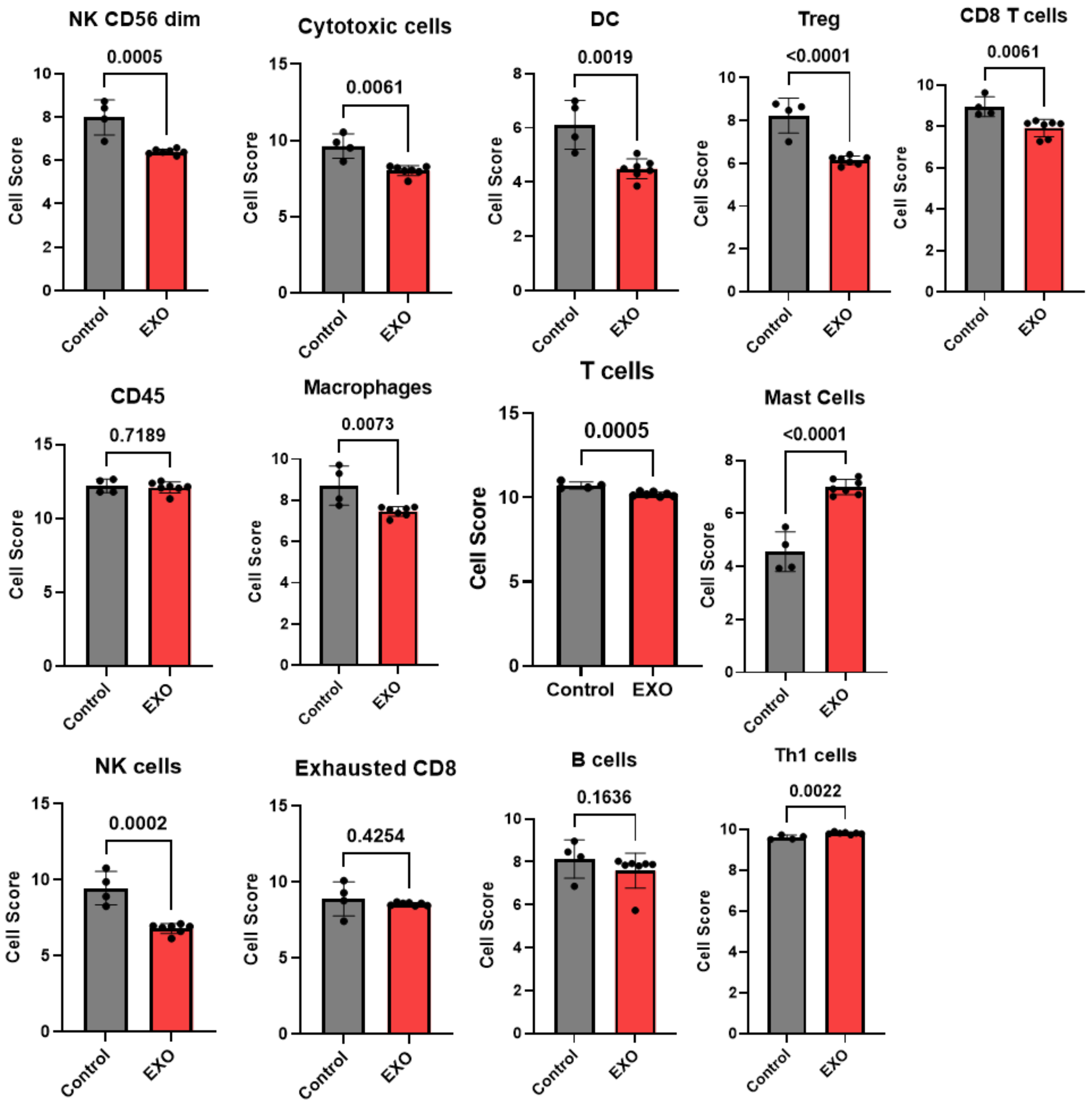


**Fig. S2 Immune Cell Type Scores in PBMCs in the absence or presence of HNSCC exosomes** Cell type scores from NanoString transcriptomic data showing immune cell abundance in PBMCs in the absence (Control) or presence of HNSCC exosomes (EXO). from untreated healthy donors (n=4; Control) and from healthy donors treated with exosomes (n=7; EXO) at a concentration of $1.04\times10^{9}$ particles/ml. The EXO group included PBMCs from four healthy donors treated with exosomes derived from three different HNSCC patients (HNC208, HNC285, and HNC365); three of the donors were each treated with exosomes from two different patients. Significance between groups was determined by either Mann-Whitney rank sum test (cytotoxic cells, B cells and CD8 T cells) or unpaired t-test. Bars represent means ± SD, symbols represent individual treatment conditions. The abundance of the different immune cell types (at the RNA level) in the various patient cohorts was calculated as log2 cell type scores. The cell scores for a specific cell type can only be compared between two groups (control vs EXO) but do not support claims that a cell type is more abundant than another cell type within the same group.

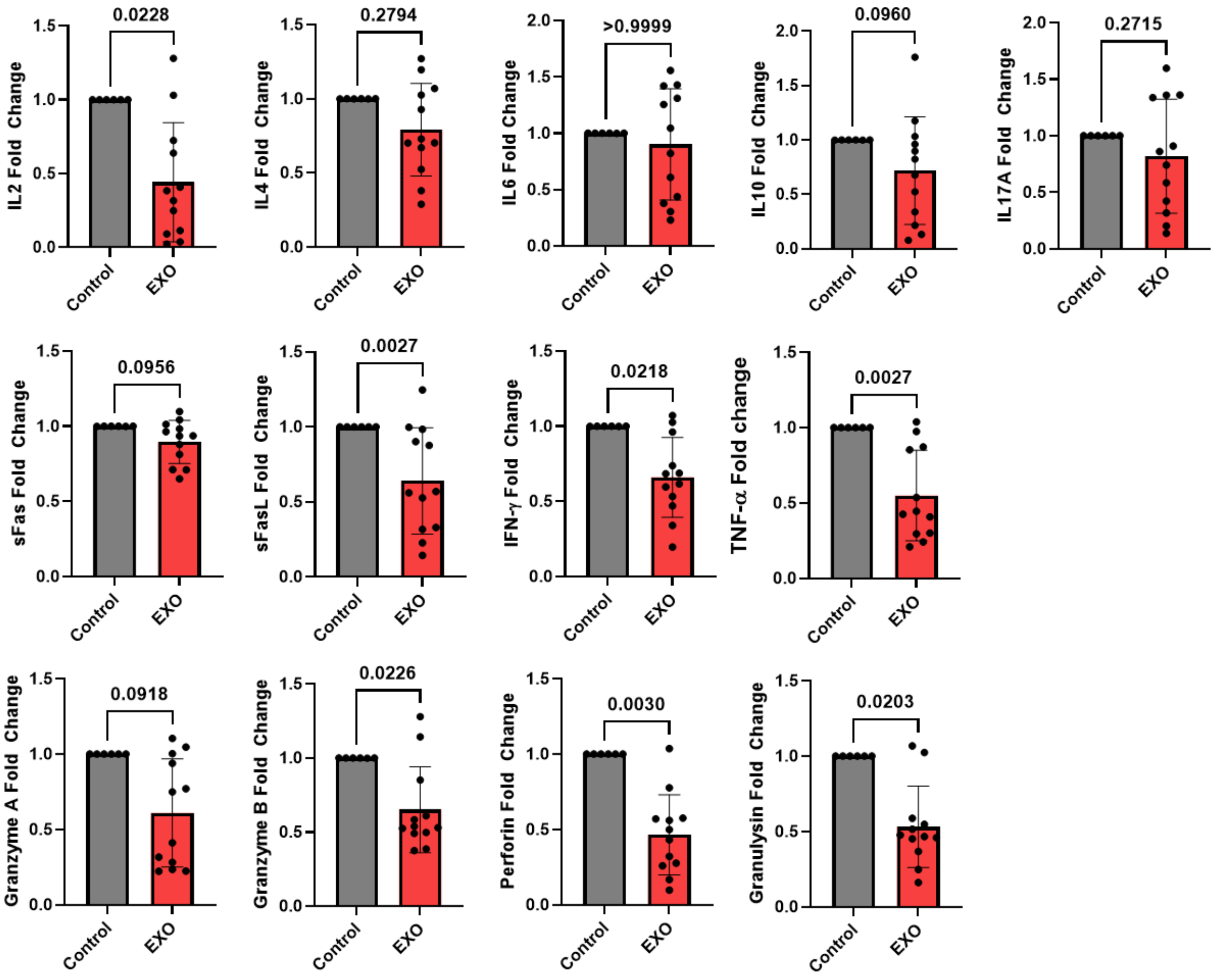


**Fig. S3 Cytokine and cytotoxicity marker release profiles of activated CD8⁺ T cells in the absence or presence of HNSCC exosomes.** Multiplex cytokine release assay showing Fold change in the abundance of individual proteins in activated $CD8^+$ T cells that were treated with exosomes from HNC208, 285 and 365 (EXO, 0.5 X $10^9$ particles/ ml) as compared to untreated controls. Data were normalized to protein levels in controls. Data were measured in n = 6 healthy donors (the same donors were used for both treatments). Statistical significance was determined by Mann-Whitney rank sum test.

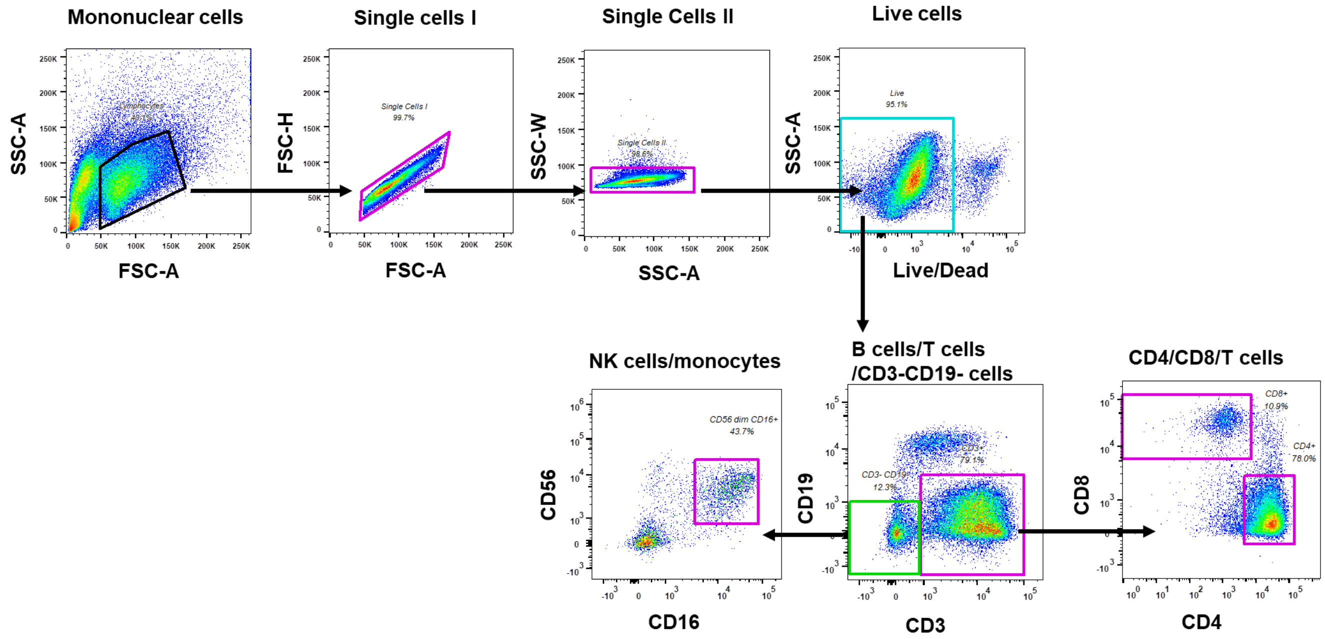


**Fig. S4. Gating strategy for immune cell phenotyping flow cytometry experiments.** Representative gating to identify various immune cell phenotypes from PBMCs from a single healthy donor is shown here. Briefly, out of the total events acquired, we first gated the mononuclear cell popular, followed by single cells and live cells (gated as Zombie UV$^{+}$). The live cell population was then gated as CD3-BV421$^{+}$ T cells, CD19-APC-Cy7$^{+}$ B cells or CD3$^{-}$CD19$^{-}$ population. The CD3$^{+}$ T cells were further classified into CD4-BV605$^{+}$ and CD8-BV711$^{+}$ subpopulations. The CD56-BV785 $^{dim}$ CD16$^{-}$ NK cell population was identified within the CD3-CD19$^{-}$ population. We then subdivided the monocyte subsets in the CD3$^{-}$CD19$^{-}$ population based on the expression of CD16-APC and CD14-BV510. In all of the immune cell subsets that were defined as shown, we then measured the Kv1.3 (Alexa Fluor 555) and KCa3.1 (ATTO 488) abundance.

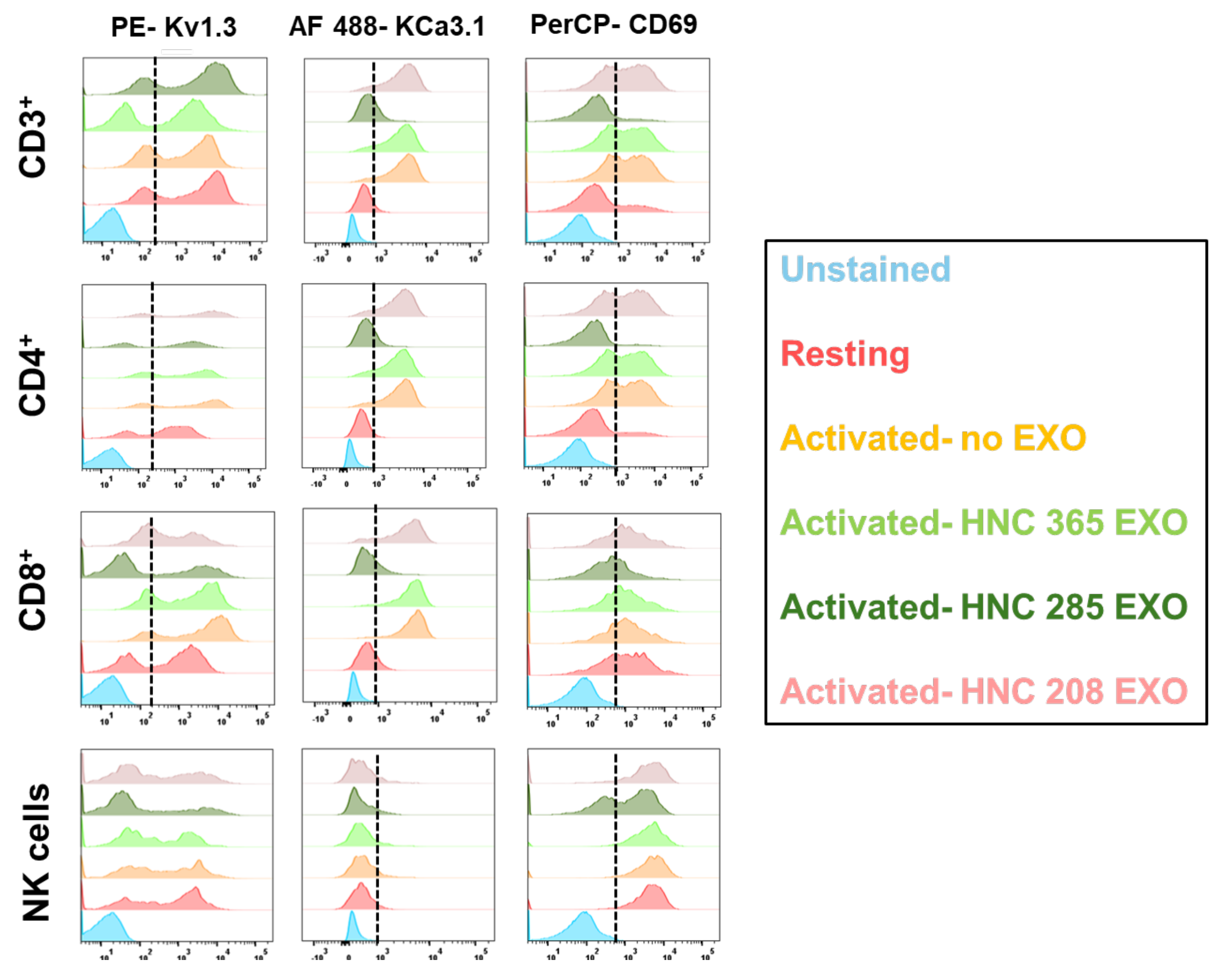


**Fig. S5 Kv1.3, KCa3.1, CD69 expression in immune cell subsets in the absence or presence of HNSCC exosomes.** Staggered histograms showing the abundance of Kv1.3, KCa3.1 and CD69 in an activated healthy donor PBMCs in the absence or presence of 1.04 X $10^9$ particles/ml HNSCC exosomes (EXO, derived from HNC208, HNC285 and HNC365 cells). Unstained cells and resting PBMCs were used as controls. Data from a representative healthy donor are shown.

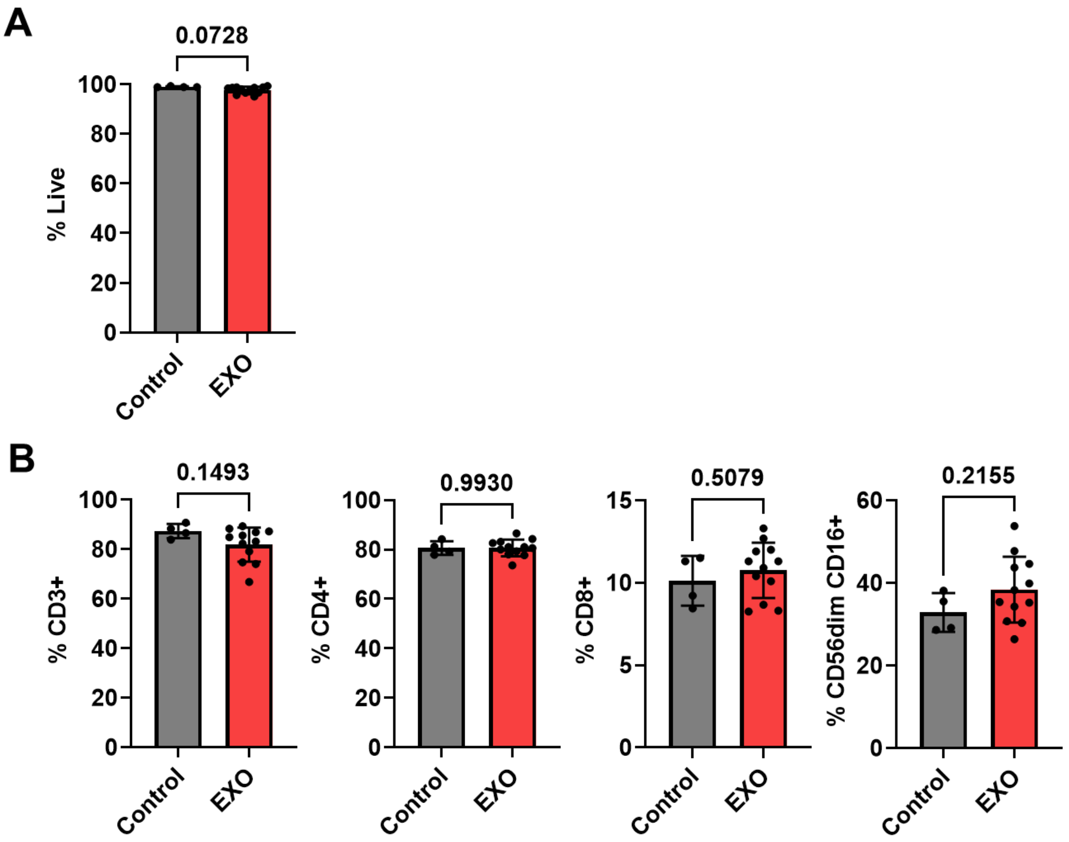


**Fig. S6 Effect of HNSCC exosomes on PBMC viability and immune cell subset abundance** (A) Cell viability, measured by flow cytometry using Zombie UV live- dead stain and (B) Quantification of abundance of various immune cell subsets in four healthy donor PBMCs in the absence (control) or presence of HNC208, HNC285 and HNC365 exosomes (EXO, 1.04 X $10^9$ particles/ ml) measured by flow cytometry. Some donors were treated with exosomes from multiple HNSCC patients. Untreated PBMC were used as controls. Significance determined by unpaired t-test. Bars represent means ± SD, symbols represent individual treatment conditions.

**Table S1 Capture antibody panels to detect Immune Checkpoint proteins (Panel 1) and Tetraspanins (Panel 2) detected on surface of intact exosomes by multiplex ELISA**

| Panel 1 | | | | Panel 2 | | |
|---|---|---|---|---|---|---|
| **Spot #** | **Antibody** | **Vendor** | **Catalog #** | **Antibody** | **Vendor** | **Catalog #** |
| 1 | CD155 | R&D systems | MAB25301 | CD9 | MSD | F215M |
| 2 | CD73 | R&D systems | MAB5795 | No antibody | | |
| 3 | CD112 | R&D systems | MAB2229 | CD63 | MSD | F215L |
| 4 | CD113 | R&D systems | MAB3064 | No antibody | | |
| 5 | No antibody | | | No antibody | | |
| 6 | No antibody | | | No antibody | | |
| 7 | No antibody | | | No antibody | | |
| 8 | CD39 | R&D systems | MAB4397 | No antibody | | |
| 9 | PDL2 | R&D systems | MAB12241 | No antibody | | |
| 10 | PDL1 | MSD | B22Z7-2 | CD81 | MSD | F215N |

**Table S2: Marker genes for individual immune cells (NanoString cell type profiling).**

| Cell Type | Gene Name |
|---|---|
| B cells | BLK |
| | CD19 |
| | MS4A1 |
| | TNFRSF17 |
| CD45 | PTPRC |
| CD8 T cells | CD8A |
| | CD8B |
| Cytotoxic cells | CTSW |
| | GNLY |
| | GZMA |
| | GZMB |
| | GZMH |
| | KLRB1 |
| | KLRD1 |
| | KLRK1 |
| | PRF1 |
| DC | CCL13 |
| | CD209 |
| | HSD11B1 |
| Exhausted CD8 | CD244 |
| | EOMES |
| | LAG3 |
| Macrophages | CD163 |
| | CD68 |
| | CD84 |

| | |
|---|---|
| Mast cells | MS4A2 |
| | TPSAB1 |
| Neutrophils | CSF3R |
| | FCGR3A |
| | S100A12 |
| NK CD56$^{dim}$ cells | IL21R |
| | KIR_Inhibiting_Subgroup_2 |
| | KIR3DL1 |
| | KIR3DL2 |
| NK cells | NCR1 |
| | XCL2 |
| T cells | CD3D |
| | CD3E |
| | CD3G |
| | CD6 |
| | SH2D1A |
| Th1 cells | TBX21 |
| Treg | FOXP3 |

**Table S3: Antibodies used for Flow Cytometry experiments**

| Marker | Fluorophore | Clone | Vendor | Catalog # | RRID |
|---|---|---|---|---|---|
| Kv1.3 | N/A | N/A | Alomone | APC-101-GP | AB_2340958 |
| IgG | Alexa-Fluor 555 | N/A | ThermoFisher | A-21435 | AB_2535856 |
| KCa3.1 | ATTO 488 | 6C1 | Alomone | ALM-051 | AB_10918529 |
| CD19 | APC-CY7 | HIB19 | Biolegend | 302218 | AB_314248 |
| CD3 | BV 421 | UCHT1 | Biolegend | 300434 | AB_10962690 |
| CD4 | BV 605 | OKT4 | Biolegend | 317438 | AB_11218995 |
| CD8 | BV 711 | SK1 | Biolegend | 344734 | AB_2565243 |
| CD56 | BV 785 | HCD56 | Biolegend | 318348 | AB_2563564 |
| CD16 | APC | 3G8 | Biolegend | 302019 | AB_492974 |
| CD14 | BV 510 | 63D3 | Biolegend | 367124 | AB_2716229 |
| CD69 | PerCP Cy 5.5 | FN50 | Biolegend | 310925 | AB_2074957 |
| Calmodulin | Alexa Fluor 647 | 2D1 | Novus Biologicals | NB120-2860 | - |
| Zombie UV | UV | N/A | Biolegend | 423107 | N/A |

**Table S4 List of differentially expressed genes**

| | Gene name | Gene family name | Immune Response Category | Annotation | p-value |
|---|---|---|---|---|---|
| 1 | AICDA | Apolipoprotein B mRNA editing enzyme catalytic subunits | T-Cell Functions | B-cell differentiation | 6.33E-12 |
| 2 | TPSAB1 | | Cell Functions | Basic cell functions, Cell Type specific | 2.11E-10 |
| 3 | CD14 | Scavenger receptors\| CD molecules | | CD molecules, Innate immune response | 3.05E-10 |
| 4 | FUT7 | Fucosyltransferases | Leukocyte Functions | Leukocyte migration | 6.81E-10 |
| 5 | IL1A | Endogenous ligands\| Interleukins | Cytokines, Interleukins | Acute-phase response, Cytokines and receptors, Inflammatory response, Innate immune | 2.48E-09 |

| | | | | | |
|---|---|---|---|---|---|
| | | | | response, Interleukins | |
| 6 | IL6 | Interleukins\| Interferons\| Interleukin 6 cytokine family | Interleukins | Humoral immune response, Interleukins | 1.27E-08 |
| 7 | TNFSF14 | Tumor necrosis factor superfamily\| CD molecules | Cytokines, Regulation, T-Cell Functions, TNF Superfamily | CD molecules, Cytokines and receptors, Regulators of T-cell activation, TNF superfamily members and their receptors, T-cell regulators, T-cell proliferation | 3.81E-08 |
| 8 | IL3 | Interleukins | Regulation, T-Cell Functions | Interleukins, Regulators of Th1 and Th2 development | 1.02E-07 |

| | | | | | |
|---|---|---|---|---|---|
| 9 | APOE | Apolipoproteins | Transporter Functions | Cell Type specific, Lipid transporter activity | 1.24E-07 |
| 10 | CXCL3 | Endogenous ligands\| Chemokine ligands | Chemokines, Regulation | Chemokines and receptors, Regulation of inflammatory response | 1.56E-07 |
| 11 | IL23A | Interleukins | Interleukins | Innate immune response, Interleukins | 2.24E-07 |
| 12 | HLA-DQA1 | C1-set domain containing\| Histocompatibility complex | Antigen Processing | Adaptive immune response, Antigen processing and presentation | 4.04E-07 |
| 13 | SLC11A1 | Solute carriers | Macrophage Functions | T-cell proliferation | 4.65E-07 |
| 14 | CXCL2 | Endogenous ligands\| Chemokine ligands | Chemokines, Regulation | Regulation of inflammatory response, Chemokines and receptors | 4.92E-07 |

| | | | | | |
|---|---|---|---|---|---|
| 15 | CREB5 | Basic leucine zipper proteins | | Complement pathway, Inflammatory response | 6.67E-07 |
| 16 | CX3CR1 | C-X-3-C motif chemokine receptors | Chemokines, Microglial Functions | Adaptive immune response, Chemokines and receptors, Microglial cell activation | 7.16E-07 |
| 17 | SH2D1B | SH2 domain containing | Leukocyte Functions | Leukocyte activation | 1.55E-06 |
| 18 | SERPINB2 | Serpin peptidase inhibitors | Senescence | Senescence initiators interferon related | 1.79E-06 |
| 19 | THBD | CD molecules\| C-type lectin domain containing | Leukocyte Functions | CD molecules, Leukocyte migration | 2.17E-06 |
| 20 | LILRB2 | Inhibitory leukocyte immunoglobulin like receptors\| CD molecules\| Ig-like cell adhesion molecule family | Regulation | CD molecules, Regulation of immune response | 4.33E-06 |

| | | | | | |
|---|---|---|---|---|---|
| 21 | CCL18 | Chemokine ligands | Chemokines | Chemokines and receptors, Anti-inflammatory cytokines | 9.50E-06 |
| 22 | CD276 | C2-set domain containing\| CD molecules\| V-set domain containing | Regulation | CD molecules, Regulation of immune response | 1.13E-05 |
| 23 | IL9 | Interleukins | Cytokines | Cytokines and receptors, Interleukins, Regulation of inflammatory response | 2.43E-05 |
| 24 | TLR4 | CD molecules\| Toll like receptors | Microglial Functions, TLR | CD molecules, Innate immune response, Microglial cell activation, Toll-like receptor | 3.40E-05 |

| | | | | | |
|---|---|---|---|---|---|
| 25 | KLRG1 | Killer cell lectin like receptors\| C-type lectin domain containing | NK Cell Functions, Regulation | Innate immune response, NK cell functions, Regulation of immune response | 3.99E-05 |
| 26 | LILRB3 | Inhibitory leukocyte immunoglobulin like receptors\| CD molecules | Regulation | CD molecules, Regulation of immune response | 5.12E-05 |
| 27 | PLAU | | Senescence | Senescence pathway | 5.51E-05 |
| 28 | NLRP3 | NLR family\| Pyrin domain containing | | Innate immune response | 6.43E-05 |
| 29 | IL23R | | Cytokines | Cytokines and receptors | 6.85E-05 |
| 30 | CCL8 | Endogenous ligands\| Chemokine ligands | Chemokines, Regulation | Chemokines and receptors, Regulation of inflammatory response | 6.98E-05 |

| | | | | | |
|---|---|---|---|---|---|
| 31 | MSR1 | Scavenger receptors\| Scavenger receptor cysteine rich domain containing\| CD molecules | Cell Functions | Basic cell functions, Cell Type specific, CD molecules | 7.54E-05 |
| 32 | CCL20 | Endogenous ligands\| Chemokine ligands | Chemokines | Chemokines and receptors | 8.46E-05 |
| 33 | MRC1 | Scavenger receptors\| CD molecules\| C-type lectin domain containing | | CD molecules | 0.00016607 |
| 34 | THBS1 | | Antigen Processing, Cell Cycle, Regulation | Cell cycle arrest, Cell Type specific, Chronic inflammatory response, Negative regulation of antigen processing | 0.00017981 |
| 35 | LILRA1 | Activating leukocyte immunoglobulin like receptors\| CD molecules | Regulation | CD molecules, Regulation of immune response | 0.00018715 |
| 36 | AXL | Immunoglobulin like domain containing\| V-set domain containing\| Fibronectin type III | | Innate immune response | 0.00023864 |

| | | | | | |
|---|---|---|---|---|---|
| | | domain containing\| Receptor Tyrosine Kinases | | | |
| 37 | NRP1 | CD molecules | Cell Functions | Basic cell functions, CD molecules | 0.00025617 |
| 38 | CD36 | Scavenger receptors\| CD molecules | Transporter Functions | CD molecules, Receptors involved in phagocytosis | 0.00039933 |
| 39 | CX3CL1 | Endogenous ligands\| Chemokine ligands | Chemokines, Leukocyte Functions | Adaptive immune response, Leukocyte activation, Chemokines and receptors, Th1 orientation | 0.00042197 |
| 40 | CMKLR1 | Chemerin receptor | Chemokines | Chemokines and receptors | 0.00043841 |
| 41 | IL11 | Endogenous ligands\| Interleukins\| Interleukin 6 cytokine family | B-Cell Functions, Cytokines, Interleukins, | Anti-inflammatory cytokines, B-cell | 0.00049627 |

| | | | | | |
|---|---|---|---|---|---|
| | | | T-Cell Functions | differentiation, Interleukins | |
| 42 | TLR8 | CD molecules\| Toll like receptors | TLR | CD molecules, Innate immune response, Toll-like receptor | 0.00051752 |
| 43 | FOXJ1 | Forkhead boxes | | Cell Type specific, Humoral immune response | 0.00055677 |
| 44 | CD160 | Immunoglobulin like domain containing\| CD molecules | Regulation | Cell Type specific, CD molecules, Regulation of immune response | 0.00066123 |
| 45 | ITGB3 | Integrin beta subunits\| CD molecules | Adhesion | Adhesion, CD molecules | 0.00075606 |
| 46 | CLEC7A | Scavenger receptors\| CD molecules\| C-type lectin domain containing | | Innate immune response | 0.00092245 |
| 47 | HLA-DQB1 | C1-set domain containing\| Histocompatibility complex | Antigen Processing | Adaptive immune response, | 0.00121075 |

| | | | | | |
|---|---|---|---|---|---|
| | | | | Antigen processing and presentation | |
| 48 | BST1 | CD molecules | | CD molecules, Humoral immune response | 0.00129477 |
| 49 | C1QA | Complement system | Complement | Complement pathway, Innate immune response | 0.00150679 |
| 50 | CFI | Scavenger receptor cysteine rich domain containing\| Complement system | | Innate immune response | 0.00156695 |
| 51 | MARCO | Scavenger receptors\| Scavenger receptor cysteine rich domain containing | | Cell Type specific, Innate immune response | 0.00169533 |
| 52 | KIR_Activating_Subgroup_1 | | NK Cell Functions, Regulation | CD molecules, NK cell functions, Regulation of | 0.00195088 |

| | | | | | |
|---|---|---|---|---|---|
| | | | | immune response | |
| 53 | EOMES | T-boxes | T-Cell Functions | CD8-positive, T-cell differentiation | 0.00239442 |
| 54 | AMBP | Lipocalins | Regulation | Negative regulation of immune response | 0.0031724 |
| 55 | CD163 | Scavenger receptors\| Scavenger receptor cysteine rich domain containing\| CD molecules | Transporter Functions | CD molecules, Phagocytosis | 0.00329129 |
| 56 | CLEC5A | C-type lectin domain containing | | Innate immune response | 0.00550031 |
| 57 | CSF1R | Immunoglobulin like domain containing\| CD molecules\| Receptor Tyrosine Kinases | | CD molecules, Innate immune response | 0.01109937 |
| 58 | TREM1 | CD molecules\| V-set domain containing | | CD molecules, Humoral immune response | 0.01151486 |

| | | | | | |
|---|---|---|---|---|---|
| 59 | CXCR2 | CD molecules\| C-X-C motif chemokine receptors\| Interleukin receptors | Chemokines, Regulation | CD molecules, Chemokines and receptors, Regulation of inflammatory response | 0.01255581 |
| 60 | CD209 | Scavenger receptors\| CD molecules\| C-type lectin domain containing | DC | Basic cell functions, Cell Type specific, CD molecules | 0.0164046 |
| 61 | MPPED1 | | Cell Functions | Basic cell functions, Cell Type specific | 0.02375641 |
| 62 | IL34 | Interleukins | Interleukins | Innate immune response, Interleukins | 0.02628454 |
| 63 | CCR9 | CD molecules\| C-C motif chemokine receptors | | CD molecules, Innate immune response | 0.03552989 |
